# Investigating microglia-neuron crosstalk by characterizing microglial contamination in human and mouse Patch-seq datasets

**DOI:** 10.1101/2022.09.07.507009

**Authors:** Keon Arbabi, Yiyue Jiang, Derek Howard, Anukrati Nigam, Wataru Inoue, Guillermo Gonzalez-Burgos, Daniel Felsky, Shreejoy J. Tripathy

## Abstract

Microglia are dynamic immune cells with diverse functional roles, including the regulation of neuronal excitability. Here, we leveraged an inconvenient truth of neuronal Patch-seq datasets — that they routinely display evidence of contamination by surrounding microglia — to better understand aspects of microglia-neuronal crosstalk. We first quantified the presence of microglial transcripts in three Patch-seq datasets of human and mouse neocortical neurons and observed extensive off-target contamination by microglia in each. Variation in microglial contamination was explained foremost by donor identity, especially in human samples, and neuronal cell type identity. Differential expression testing and enrichment analyses suggest that microglial contamination in Patch-seq is reflective of activated microglia and that these transcriptional signatures are distinct from those captured via single-nucleus RNAseq. Finally, neurons with greater microglial contamination differed markedly in their electrophysiological characteristics, including lowered input resistances and more depolarized action potential thresholds. Our results suggest microglial contamination contributes to cell- and donor-related electrophysiological variability and sheds light on how microglia might impact neurons *in vivo*.

## Introduction

Microglia are the innate immune cells of the central nervous system and exhibit a complex array of phenotypes and functions (Bachiller et al., 2018; Ginhoux et al., 2013). Microglia frequently interact with neurons and have well-established roles in sculpting synaptic connections and regulating their plasticity (Cornell et al., 2022). For example, pruning synapses is crucial for the refinement of neuronal circuitry during development as well as learning and memory in the adult brain. Synaptic pruning is regulated by phagocytic microglia and by several immune-related signaling pathways (Lehrman et al., 2018; Sellgren et al., 2017; Wang et al., 2020), and has important emerging causal links to neuropsychiatric disease (Parellada and Gassó, 2021; Sellgren et al., 2019). The strength of retained functional synapses are modified by microglia through the release of various cytokines (Basilico et al., 2019; Hoshino et al., 2017; Kim et al., 2020; Lin et al., 2019; Pribiag and Stellwagen, 2014; Sun et al., 2020).

There is growing evidence that microglia function beyond the synapse to regulate aspects of intrinsic neuronal excitability. Microglia actively survey neural circuit excitability and can undertake a role similar to inhibitory cells to maintain the synchrony and homeostasis of local neuron activity (Badimon et al., 2020). Microglia achieve this by forming direct contacts with neuronal cell bodies (Akiyoshi et al., 2018; Cserép et al., 2020) or by secreting specific molecules such as ATP, TNF-α, IL1B, and other cytokines (Hikosaka et al., 2022). However, the effects of this regulation are highly varied among different neuronal types and brain regions. For example, microglial activation decreases intrinsic excitability among neocortical pyramidal cells (Yamawaki et al., 2022) but increases excitability among cerebellar purkinje cells via phosphatase-dependent signaling and SK channel regulation (Yamamoto et al., 2019). Microglia further differ in their transcriptomic states across neocortical layers, with differences in the densities of homeostatic and proliferative microglial subtypes in upper relative to lower layers (Stogsdill et al., 2022). Despite the importance of microglia for neuronal function, much of this work has been done using rodent models, and it is unclear how such effects might translate to humans or generalize across diverse neuronal types.

Patch-seq is a recently developed method for characterizing multi-modal neuronal diversity by combining patch-clamp electrophysiology with single-cell RNA-sequencing (scRNA-seq) (Lipovsek et al., 2021). Unlike more traditional methods for scRNA-seq that first rely on cell dissociation, Patch-seq is unique in that the patch pipette is used to carefully harvest mRNA from the cytoplasm and nucleus of the targeted neuron following electrophysiology (Cadwell et al., 2016; Lipovsek et al., 2021). A major challenge with Patch-seq, however, is that off-target mRNA contamination from surrounding cells in the local microenvironment is often prevalent in these datasets. We initially reported the paradoxical presence of high levels of gene expression markers of non-neuronal cells (Tripathy et al., 2018), including microglia, in multiple Patch-seq datasets of confirmed neurons (Cadwell et al., 2015; Földy et al., 2016; Fuzik et al., 2016). Because this contamination was only observed in datasets collected from acute brain slices but not those from sparsely plated cell cultures (Bardy et al., 2016), we reasoned that such contamination is likely due to the physical contact of the patch pipette with processes of other cells in the local microenvironment of the recorded neuron. Since our initial report, a number of Patch-seq studies have corroborated these findings, and in particular, that microglial transcripts appear especially prevalent among neuronal Patch-seq datasets (Berg et al., 2021; Lee et al., 2021a; Scala et al., 2020). However, it remains unknown whether contamination from surrounding microglia is associated with functional characteristics of the recorded neurons.

Here, we explore the potential impact of microglia on neuronal function by capitalizing on our earlier observation that contamination from surrounding microglia is widely present among neuronal Patch-seq datasets. In particular, we survey the magnitude and transcriptomic signature of microglial contamination among high-quality datasets of Patch-seq profiled neurons from human and mice acute brain slices. We further asked if there are particular aspects of donor, neuronal type, and other technical and biological factors that are most associated with microglial contamination. Critically, we aimed to learn if neurons with a high degree of contamination from surrounding microglia also tend to have altered intrinsic electrophysiological characteristics. Our findings indicate that microglial contamination in Patch-seq is reflective of a distinct activated and infiltrating microglial state and is further associated with profound alterations in neuronal electrophysiology.

## Methods

### Datasets

All datasets were obtained from the Allen Brain Institute Cell Types Database (http://celltypes.brain-map.org/) or the Tolias/Berens/Sandberg labs (https://github.com/berenslab/mini-atlas).

### Patch-seq

We made use of data from three human and mouse Patch-seq experiments, as detailed previously (summarized in Table 1 and described below).

**Table 1.** Description of Patch-seq datasets used in this study.

| Dataset Reference | Species | Brain Region | Cell types sampled | N cells | N donors/ animals |
| --- | --- | --- | --- | --- | --- |
| Berg et al., 2021 | Human | Medial Temporal Gyrus | Superficial pyramidal cells | 278 | 48 |
| Gouwens et al., 2020 | Mouse | Visual cortex | GABAergic cells | 4,270 | 1,040 |
| Scala et al., 2020 | Mouse | Motor cortex | Glutamatergic and GABAergic cells | 1329 | 266 |

For data from Allen Brain Institute human Patch-seq experiments (Berg et al., 2021), briefly, surgical tissues were obtained from local Seattle-area hospitals and transported in NMDG-based artificial cerebral spinal fluid (ACSF) to a laboratory, where acute slices were prepared for patch clamp recording. Electrophysiological recordings were performed in warm (32–34 °C) recording ACSF in the presence of pharmacological blockers of fast glutamatergic and GABAergic synaptic transmission. Upon completion of the electrophysiology experiment, negative pressure was applied to the patch pipette to extract the cytosol and nucleus of the target cell (Figure 1B). The SMART-Seq v4 platform was used for cDNA amplification and library construction, followed by sequence alignment. Single-cell transcriptomes from Patch-seq samples were subsequently mapped to a reference taxonomy based on single-nucleus RNAseq data from the human medial temporal gyrus (Berg et al., 2021; Hodge et al., 2019). RNA-sequencing metadata and counts data were downloaded for our analyses (accessed 05/10/2021). A total of 278 supragranular glutamatergic neurons from MTG in 48 subjects (18 female) were included. Brain slices not used for slice electrophysiology were also characterized using immunohistochemistry for various protein markers, including IBA1, marking microglia, and GFAP, marking astrocytes.

**Figure 1.**
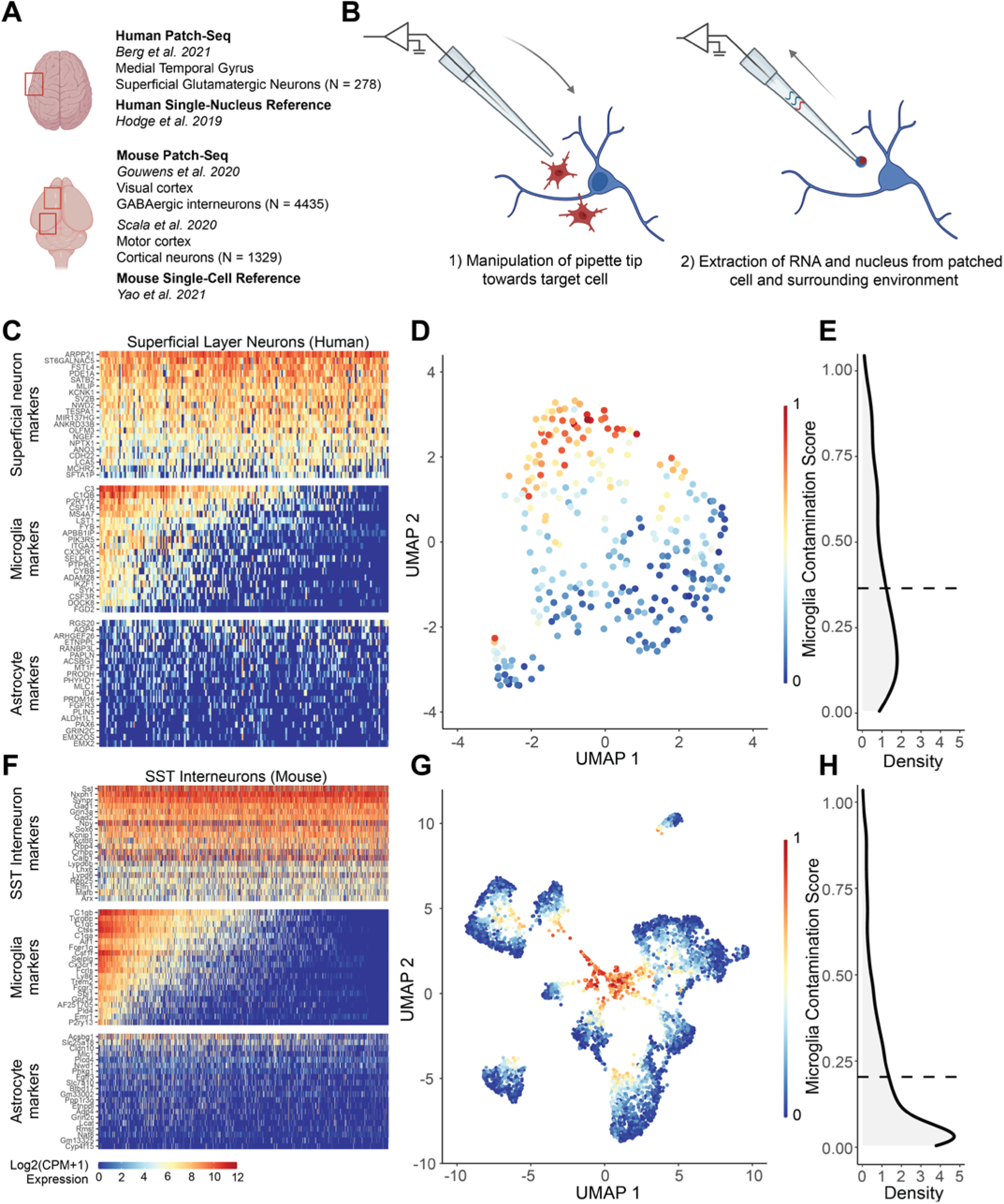
Patch-seq transcriptomes of human and mouse neurons express off-target expression of microglial marker genes at levels sufficient to drive unbiased clustering. **(A)** Overview of datasets used in the current study, including three recent Patch-seq datasets of cortical neurons in human and mouse, with comparisons made to dissociated human single nucleus and mouse single cell RNA-sequencing datasets, respectively. **(B)** Schematic illustrating manipulation of patch-pipette towards a neuron of interest, and collection of mRNA from cell nucleus, cytoplasm, and possibly the surrounding environment via the patch-pipette. **(C**,**F)** Gene expression profiles for human superficial glutamatergic neurons (C, Berg dataset) or mouse SST interneurons (F, Gouwens dataset.) for various cell type-specific markers. Each row represents a cell type-specific marker gene and columns represent individual neurons, ordered from left to right by decreasing microglial contamination score. **(D**,**G)** Low-dimensional visualization of transcriptomes from neuronal Patch-seq samples clustered by most variable gene expression and color-coded by microglial contamination score for human pyramidal cells (D, Berg dataset), and mouse GABAergic interneurons (G, Gouwens dataset). **(E**,**H)** Distribution density of microglial contamination scores for human (E) and mouse (H) neurons, dashed lines indicate population mean.

We also made use of data from Allen Brain Institute mouse Patch-seq experiments as detailed previously (Gouwens et al., 2020). Animals were anesthetized using isoflurane and intracardially perfused with ice-cold NMDG-based ACSF prior to slice preparation. Cell recordings were performed using male and female mice between the ages of P45 and P70. Multiple Cre-driver lines were used to target cells for morphoelectric and transcriptomic characterization. Electrophysiological recordings were performed in warm (34 °C) recording ACSF in the presence of pharmacological blockers of fast glutamatergic and GABAergic synaptic transmission. The SMART-Seq v4 platform was used for cDNA amplification and library construction, followed by sequence alignment. Single-cell transcriptomes from Patch-seq samples were then mapped to a reference taxonomy of dissociated neuronal cells from mouse VISp (Tasic et al., 2018). RNA-sequencing metadata and counts data were downloaded for further analyses (v2, released 07/01/2020). A total of 4,270 GABAergic interneurons sampled from visual cortex in 1,040 mice (468 female) were included.

We also made use of mouse data from Scala et al, reflecting mouse Patch-seq experiments (Scala et al., 2020). Animals were anesthetized using isoflurane and decapitated prior to rapid brain removal and collection into cold NMDG-based ACSF for slice preparation. Cell recordings were performed using male and female mice between the ages of P35 and P245. Multiple Cre-driver lines were used to target cells for morphoelectric and transcriptomic characterization. Electrophysiological recordings were performed at room temperature (25 °C) using recording ACSF where fast synaptic transmission was not blocked. The Smart-seq2 protocol was used for cDNA amplification and library construction, followed by sequence alignment. Single-cell transcriptomes from patch-seq samples were mapped to multiple taxonomies, including a taxonomy based on cells from mouse VISp and anterolateral motor cortex (ALM), enabling direct comparison with cells from (Gouwens et al., 2020). RNA-sequencing metadata and counts data were downloaded for further analyses (accessed 08/01/2022). A total of 1329 glutamatergic and GABAergic neurons sampled from the motor cortex in 266 mice (135 female) were included.

### Single-cell and single-nucleus RNA-seq derived from dissociated cells and nuclei

Gene expression patterns from Patch-seq samples were compared to cell dissociation-based single-cell and single-nucleus RNA-seq datasets. Hodge et al. (2019) used snRNA-seq and multiple clustering approaches to describe the cell type taxonomy of 15,928 nuclei from human MTG. Yao et al. (2021) used scRNA-seq to profile ∼1.3 million cells from the adult mouse isocortex and hippocampal formation to generate a transcriptomic cell-type taxonomy. Cells and nuclei transcriptomes quantified with SMART-seq were downloaded from both datasets and used for the present analyses (accessed 09/25/2021).

### Microglial contamination score

To quantify the extent of microglial contamination for each Patch-seq cell, we applied previously described methods for assessing the quality of Patch-seq transcriptomic data (Lee et al., 2021b; Tripathy et al., 2018). Specifically, we employed cell type-specific markers for human and mouse neocortical neurons calculated in Lee et al. (2021) (https://github.com/AllenInstitute/patchseqtools). These markers describe both “on” markers, or genes that are highly and ubiquitously expressed in neurons of interest relative to other cell types. “Off” markers are expected to be expressed at low levels in Patch-seq cells, and if expressed together with “on” markers, are an indicator of possible cellular contamination. Lee et al. calculated both types of markers using the single-cell and single-nucleus RNA-seq datasets from dissociated cells and nuclei described previously (Hodge et al., 2019; Tasic et al., 2018), which can serve as ground truth expression data for comparison with Patch-seq cells (Lee et al., 2021b).

Using the definitions described in (Lee et al., 2021b; Tripathy et al., 2018), we calculated the *contamination score* of microglia *(M)* in a neuron subtype of interest *(N)*, as:

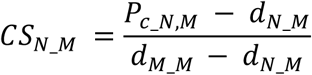

The expression of microglia markers in a Patch-seq cell *c* are compared to the expected expression in reference data from which the markers were derived. To do this we take the summed normalized expression (log2 CPM) of 50 microglia markers in a cell of *N (P*_*c_N, M*_) and subtract the median microglia marker expression in dissociated cells of type *N (d*_*N_M*_). If this numerator is negative (for example, if cell *c* expresses no microglial markers but *d*_*N_M*_ is positive), it is set to 0 in these cases (indicating no detected contamination). The denominator scales this value by the expected expression of microglial markers in reference microglia. The contamination score can thus be interpreted as a ratio of the excess off-target microglial expression, scaled between 0 and 1 (where 1 indicates that the cell expresses off-target microglial expression at a level similar to actual microglia).

### Visualizing Patch-seq gene expression

Gene expression from human and mouse Patch-seq cells was processed according to the standard Seurat V3 workflow (https://satijalab.org/seurat) (Butler et al., 2018; Stuart et al., 2019). All available cells for human and mouse were included in the following steps. Best practices for processing Patch-seq data typically involve filtering out genes that are highly expressed in non-neuronal cell types, however this step was omitted to gauge the extent of off-target contamination. Only mitochondrial genes and genes of uncertain function were filtered out prior to normalization. The top 2000 variable features (5000 for mouse) were identified, followed by scaling, linear dimensionality reduction, and clustering using the default parameters. Cells were visualized by projection onto Uniform Manifold Approximation and Projection (UMAP) space.

### Associations between microglial contamination and donor, cell type, and tissue characteristics

Univariate relationships between microglia contamination score and available Patch-seq metadata were described, including neuron transcriptomic-type, cell soma depth from the pial surface (in micrometers), and, in human samples, histological markers of tissue pathology. Histological pathology scores for IBA1 and GFAP protein expression were included as the only markers not skewed heavily towards zero and were binned by marker score (low ≤ 1, high > 1) (Berg et al., 2021). Given the large number of mouse transcriptomic types at the cluster resolution, we aggregated closely related cell clusters by summarizing t-types using the subclass designation and the first marker gene (e.g., Sst Hspe Sema3c → Sst Hspe).

To further explain the influence of these characteristics with off-target microglial expression in Patch-seq samples, we employed mixed effects models. Only cells with data available for the variables of interest were included, and cell types with fewer than 20 cells in either the Berg or Gouwens datasets (or 10 cells in the Scala dataset) were excluded. The following model was used for the human Patch-seq data:

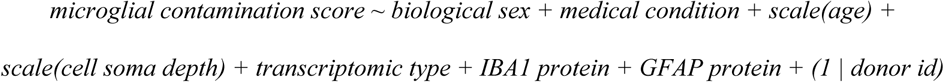

And for the Gouwens mouse Patch-seq data:

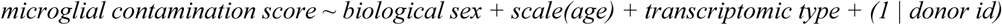

And for the Scala mouse Patch-seq data:

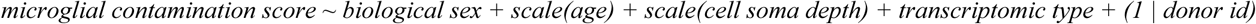

We estimated the proportion of variation in microglial off-target contamination predicted by each variable. The coefficient of determination (R^2^) for each fixed effect (every term except donor id) was iteratively calculated by taking the difference between the marginal R^2^ value of the full model and the marginal R^2^ value of a reduced model, in which the fixed effect of interest was replaced by a random intercept. Marginal R^2^ was calculated using the method described by (Nakagawa et al. 2017) and implemented in the *MuMIn* package (Burnham and Anderson, 2010). The variance explained by the random effect of donor id was estimated by subtracting the marginal R^2^ from the conditional R^2^ of the full model.

### Differential gene expression analysis

To characterize the transcriptional signature associated with off-target microglial contamination in Patch-seq data, we performed differential expression testing. For the human data, high and low contamination groups were defined by the top and bottom quartiles of Patch-seq sampled neurons ranked by contamination score, respectively. For the mouse data, deciles were used to adjust for the smaller fraction of cells with high contamination. In each dataset, differential expression tests between these groups were performed with DESeq2 (Monat et al., 2019) using the FindMarkers function with default settings in Seurat v3. Differentially expressed genes (DEGs) were defined as having >2.5 or <-2.5 log2 fold-change and p-value < 0.01. In effect, this approach identifies neuronal transcripts that are significantly co-expressed with microglial contamination.

For comparison, we also performed differential expression testing between groups of microglia and neurons in the reference single-cell and single-nucleus RNA-seq datasets. We selected neuronal transcriptomic types that most closely matched those surveyed in the human and mouse Patch-seq datasets. For the Yao dataset, cells labeled “Micro-PVM” at the subclass level in all brain regions (N = 178) and “GABAergic” at the class level in VIS and VISp (N = 6846) were compared with DESeq2 as described above. For the Hodge dataset, cells labeled “Microglia’’ at the subclass level (N = 246) and cells with cluster labels including “Exc L2-3”, “Exc L2-4”, and “Exc L3-4” in MTG (N = 3609) were compared. DEGs from these tests were selected as described above.

The functional context of DEGs was explored with gene set enrichment analysis. We include 19 unique microglial gene lists from 4 publications pulled from the literature, selected for their inclusion of homeostatic and activated microglia populations from both human and mouse. Friedman et al. performed a meta-analysis of purified mouse CNS myeloid cells across a wide range of disease models, identifying distinct modules of co-regulated genes associated with lipopolysaccharide (LPS) injection, interferon signaling, and neurodegeneration. In a single-cell dataset of a mouse model of Alzheimer’s disease (AD), each module represented distinct types and activation states of microglia (Friedman et al., 2018). Keren-Shaul et al. identified disease-associated microglia (DAM) in an AD-transgenic mouse model that potentially restricts neurodegeneration (Keren-Shaul et al., 2017). Mathys et al. confirmed the presence of DAM and identified several other disease stage-specific microglial states in a mouse model of AD-like neurodegeneration (Mathys et al., 2017). Olah et al. performed scRNA-seq to explore the population structure of live microglia purified from surgically resected human cortex and characterized nine clusters of microglia encompassing states specific to homeostasis, proliferation, response to cellular stress, and responses to injury and disease (Olah et al., 2020). These findings reflect the heterogeneity of microglia, and the presence of activation and/or disease-related subtypes. Gene sets were directly acquired from supplemental materials. If human or mouse gene symbols were not available, the provided set was converted using the getLDS() function from the biomaRt package (Durinck et al., 2005, 2009). Cluster-defining genes provided by OIah et al. were filtered to those significantly downregulated in >6 other clusters to enhance specificity.

The enrichment of these gene sets in genes significantly upregulated (>2.5 log2 fold-change, p-value < 0.01) in high microglial contamination Patch-seq cells and reference scRNA-seq microglia was determined with the hypergeometric test implemented by the HypeR package (Federico and Monti, 2020).

### Electrophysiology analyses

Raw electrophysiological traces from the human (Berg et al., 2021) and mouse (Gouwens et al., 2020) Patch-seq experiments were obtained from DANDI (dataset IDs: 000209 and 000020). Stimulation protocols consisted of long-square hyperpolarizing and depolarizing current injections. The Intrinsic Physiology Feature Extractor (IPFX) toolbox (Lee et al., 2021a) was used to extract electrophysiological features from each recorded neuron, similar to as described previously (Moradi Chameh et al., 2021). Sweeps explicitly tagged as failed were discarded prior to feature extraction. Extracted features include subthreshold features (i.e., input resistance, sag ratio), action potential properties (i.e., action potential half-width, threshold time and voltage) derived from the rheobase spike, as well as multi-action potential spike train features derived from the IPFX-defined “hero” sweep (i.e., adaptation index). Action potential amplitude was defined as the difference between peak and threshold voltage and after hyperpolarization amplitude was defined as the difference between action potential threshold and the fast action potential trough. For electrophysiological data from the Scala mouse dataset, we employed processed electrophysiological features that were provided by the original study authors and standardized these so that they were broadly consistent with those from the other datasets.

We used a statistical approach to ask how microglial contamination is associated with cell-to-cell variability in electrophysiological features, after controlling for other aspects of cellular and donor identity.

Specifically, we used a mixed effects model, where, for human cells from the Berg dataset and glutamatergic and GABAergic cells from the Scala dataset:

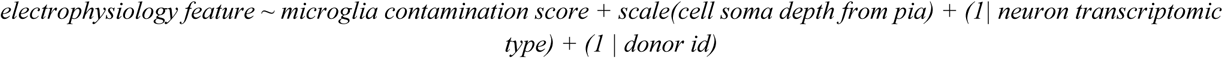

And for mouse cells from the Gouwens dataset:

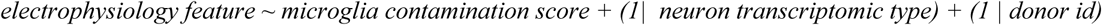

We used a log10 normalization to scale input resistance, rheobase, and action potential half-width and all electrophysiology features were standardized prior to modeling. We chose not to standardize microglia contamination scores to enable direct comparisons of effects between humans and mice, as these have very different standard deviations in microglial contamination scores. Significances for the beta coefficients associated with microglial contamination were provided using the *lmerTest* function in R (Kuznetsova et al., 2017).

## Results

### Strategy for assessing the impact of microglial off-target contamination in neuronal Patch-seq samples

To study the extent and impact of microglial off-target contamination in Patch-seq experiments we used three datasets (Figure 1A, Table 1), including 1) human supragranular glutamatergic neurons from the medial temporal gyrus previously published by the Allen Institute for Brain Sciences (Berg et al., 2021), and 2) mouse GABAergic interneurons from the primary visual cortex also previously published by the Allen Institute for Brain Sciences (Gouwens et al., 2020), and 3) mouse glutamatergic and GABAergic neurons from the primary motor cortex (Scala et al., 2020). These datasets were selected in part due to their large size, reflecting hundreds to thousands of neuronal samples. In addition, these datasets are considered to be of high-quality, reflecting thousands of measured genes per neuronal sample, as they follow extensive internal optimization procedures and strict transcriptomic and electrophysiological data quality control (Cadwell et al., 2017; Lee et al., 2021a). Lastly, the availability of high-quality cell-dissociation based single-cell and single-nucleus transcriptomes from parallel samples (Figure 1A; also collected by the Allen Institute, Hodge et al., 2019; Yao et al., 2021) enables rigorous comparisons between Patch-seq based and cell dissociation-based gene expression profiles.

### Microglial off-target contamination is widely present in human and mouse Patch-seq neuronal transcriptomes and drives unbiased clustering

To quantify the levels of microglial off-target contamination in Patch-seq-sampled neurons, we first examined the expression of cell type-specific marker genes. We observed that Patch-seq neuronal samples expressed high levels of the expected marker genes, for example, SV2B and SATB2 in human superficial pyramidal cells (Figure 1C, top row) and Sst, Gad1, and Gad2 in mouse somatostatin interneurons (Figure 1F, top row). However, further investigation revealed that many Patch-seq neuronal transcriptomes expressed high levels of multiple microglia-specific markers which are not expected to be expressed in neurons (see Methods, Lee et al., 2021a). Such markers include C3 and C1QB in human samples (Figure 1C, middle row) and C1qb and Tyrobp in mouse samples (Figure 1F, middle row). Because of the widespread upregulation of many unexpected microglia-specific transcripts in Patch-seq neuronal samples, we reason that these likely reflect the inadvertent sampling of mRNA from microglial cellular processes via the patch-pipette (Figure 1B), as opposed to the endogenous upregulation of microglia-specific transcripts by the sampled neuron. We note that the expression of astrocyte-specific markers was generally rare in both human and mouse samples (Figure 1C, 1F, bottom row), suggesting that astrocytes are likely not a major source of off-target cellular contamination in these samples. We also found qualitatively similar findings indicating the presence of multiple microglial transcripts among glutamatergic and GABAergic Patch-seq neuronal samples among the Scala dataset (Supplementary Figure 1A, B).

To obtain a single value reflective of the extent of off-target microglial contamination, we calculated microglial contamination scores for each Patch-seq neuronal cell, as defined previously in (Lee et al., 2021a; Tripathy et al., 2018). These scores provide an interpretable scalar value between 0 and 1, where 0 indicates little-to-no detected microglial contamination and 1 indicates a very high degree of contamination at levels similar to those expressed by microglia sampled via single-nucleus (human) or single-cell (mouse) RNAseq.

We asked if off-target microglial contamination is sufficient to drive unbiased transcriptional clustering of Patch-seq neuronal samples. Following standard workflows for single-cell analyses (see Methods), we found that human and mouse Patch-seq neuronal samples exhibited strong visual evidence of transcriptomic clustering, in part, according to their levels of off-target microglial contamination (Figure 1D, 1G, Supplementary Figure 1C). On average, we found that human Patch-seq neuronal transcriptomes appeared considerably more contaminated by microglia than in the mouse datasets (Figure 1E, 1H, Supplementary Figure 1D; human: 0.34 ± .25, mouse, Gouwens: 0.19 ± .22; mouse, Scala, glutamatergic: 0.15 ± 0.14; Scala, GABAergic: 0.14 ± .14, mean ± SD, normalized microglia contamination scores).

### Inter-donor differences and neuronal identity explain the most variation in microglial off-target contamination

We next wanted to understand how experimental characteristics, like cell type identity, donor characteristics, and tissue quality, might correlate with off-target microglial contamination in Patch-seq cells. Among human Patch-seq sampled superficial pyramidal neurons, we found samples collected from different neurosurgical donors varied considerably in the levels of microglial contamination (Figure 2A, Kruskal-Wallis ANOVA p = 4.3 * 10^−6^). In contrast, we found non-significant differences between transcriptomic cell identity of the sampled neuron (t-type) and microglial contamination scores (Figure 2B, Kruskal-Wallis ANOVA p = 0.22). Utilizing data from immunohistology performed on brain slices not used for electrophysiological characterization, IBA1 and GFAP protein expression (markers of microglia and astrocytes, respectively) was associated with increased microglial contamination (GFAP: Figure 2C left, Kruskal-Wallis ANOVA p = 0.067; IBA1: Figure 2C right, Kruskal-Wallis ANOVA p = 0.10). While the prior association between transcriptomically-inferred microglial contamination and IBA1 protein expression is marginal, this serves as an independent confirmation of our interpretation of increased microglial expression in samples from these donors.

**Figure 2.**
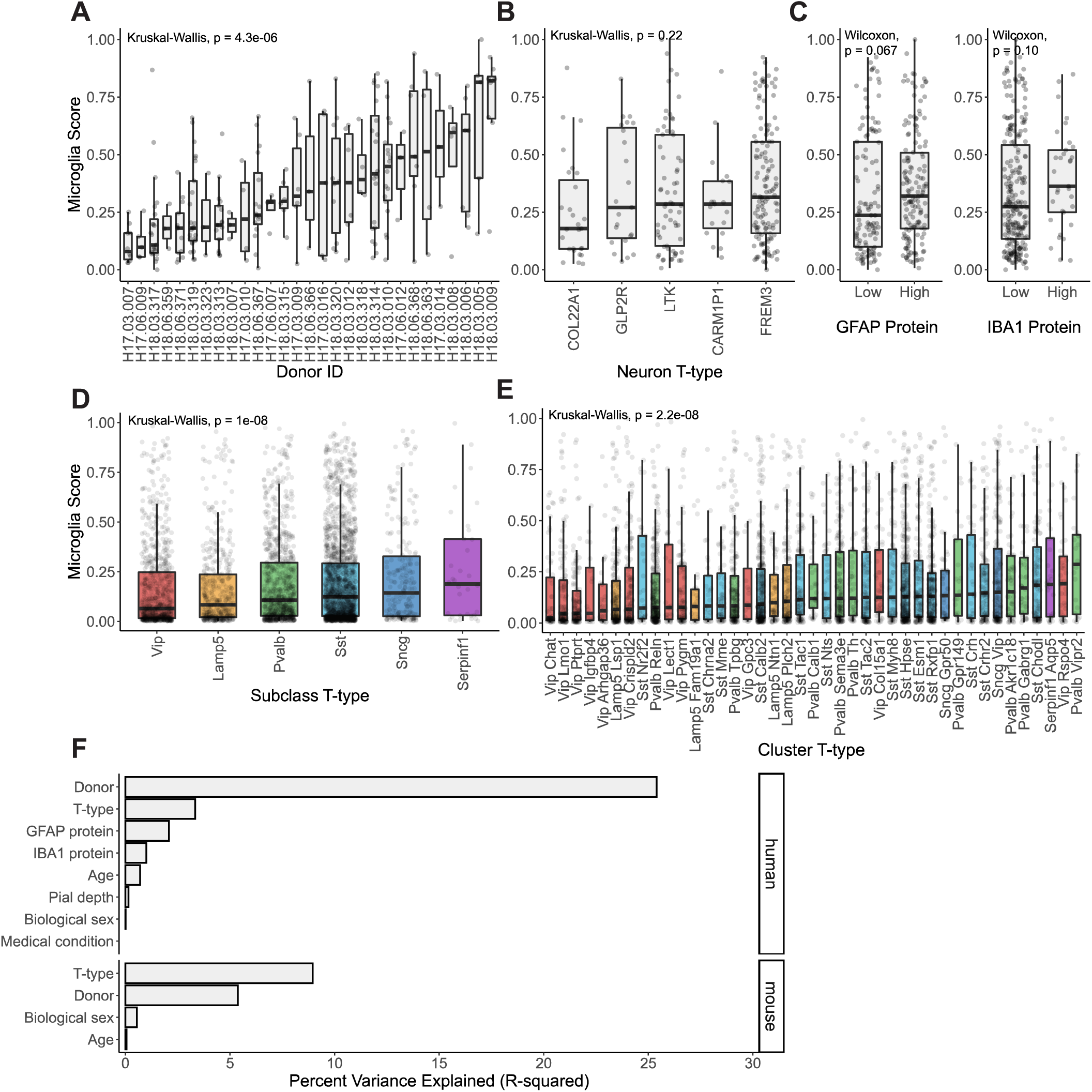
Associations between donor, cell type, and tissue characteristics with off-target microglial expression in Patch-seq samples. **(A**,**B**,**C)** Associations between microglia contamination scores (y-axis) estimated from human pyramidal neuron Patch-seq samples and neurosurgical donor identity (A), transcriptomically-inferred neuronal type identity (B), and GFAP (left) and IBA1 (right) protein expression assayed via immunohistochemistry (C) performed on brain slices not used for Patch-seq characterization (low ≤ 1, high > 1). **(D, E)** Associations between microglia contamination scores (y-axis) estimated from mouse GABAergic interneuron Patch-seq samples and interneuron cell type identity, summarized at either the subclass level (D) or cluster level (E). **(F)** Estimated percent variance explained (R-squared) in microglial contamination scores among human (top) and mouse Gouwens dataset samples (bottom) by various factors, including donor/animal identity, neuronal cell type identity (t-type), age, sex, sample depth from pial surface (Pial depth), medical condition (epilepsy or tumor), IBA1 or GFAP protein expression.

Among Patch-seq sampled GABAergic cortical interneurons from mice from the Gouwens dataset, we found that different cell types exhibited different levels of microglial contamination (subclass: Figure 2D, cluster: Figure 2E). We observed lower levels of microglial contamination among Vip and Lamp5 subclasses and comparatively higher levels in Sst, Sncg, and Serpinf subclasses (Figure 2D, Kruskal-Wallis ANOVA p = 1.0 * 10^−8^). Even among more fine-grained neuronal subtypes (i.e., cluster t-type), we observed considerably different levels of microglial contamination among clusters from the same parent subclass (Figure 2E, Kruskal-Wallis ANOVA p = 2.2 * 10^−8^). Among Patch-seq sampled GABAergic and glutamatergic neurons from the mouse Scala dataset, we also observed that transcriptomic cell type identity was also associated microglial contamination (Supplementary Figure 2B, GABAergic: Kruskal-Wallis ANOVA p = 0.097, glutamatergic: Kruskal-Wallis ANOVA p = 1.8 * 10^−12^). Moreover, we observed a modest correlation between levels of microglial contamination among matched GABAergic cell types in both the Gouwens and Scala datasets (Supplementary Figure 2C, Pearson’s R = 0.39, p =0.047). Intriguingly, both datasets showed the highest levels of microglial contamination among Pvalb Vipr2 cells (i.e., chandelier cells, (Gouwens et al., 2020), suggesting that chandelier cells might show some inherent differential vulnerability to contact by surrounding microglia (see Discussion). In addition, we found that neurons recorded deeper from the pial surface displayed less microglia contamination (Supplementary Figure 2A, GABAergic: Pearson’s R = -0.15, p = 0.00012; glutamatergic: Pearson’s R = -0.27, p = 5.4 * 10^−8^), consistent with evidence suggesting greater densities of microglia in upper relative to lower cortical layers (Stogsdill et al., 2022).

To further quantify associations between these experimental variables and microglial off-target contamination, we used a random effects model, allowing us to statistically model correlations between Patch-seq sampled neurons recorded from the same donor or animal (see Methods). Among the human pyramidal neuron samples (Figure 2G, top), donor identity, by far, explained the most neuron-to-neuron variability in microglial off-target contamination (R^2^ = 25.4%), followed by cell type identity (R^2^ = 3.34%), and other donor specific factors such as levels of GFAP and IBA1 protein expression (R^2^ = 2.08% and 1.00%, respectively). Other factors, such as donor age at time of surgery, sex, and cell soma depth from pia, were only weakly explanatory for microglial off-target contamination (R^2^ < 1%). Among mouse samples in the Gouwens dataset (Figure 2G, bottom), we found that the most important factor associated with microglial off-target contamination was cell type identity (R^2^ = 8.95%), followed by animal identity (R^2^ = 5.37%) and animal sex and age explained comparatively less variability (R^2^ = 0.553% and 0.055%). Among mouse samples in the Scala dataset, we found results qualitatively similar to those in the human dataset (Supplementary Figure 2D), with animal identity explaining the most variance, particularly in glutamatergic cells (GABAergic R^2^ = 4.99%; glutamatergic R^2^ = 15.1%), followed next by cell type identity (GABAergic R^2^ = 3.55%; glutamatergic R^2^ = 4.45%), with other factors, such as age, sex, cell soma pial depth explaining <2% of variance, each. In total, considerably more neuron-to-neuron variation in microglial off-target contamination could be explained for human samples relative to each of the mouse datasets (R^2^ = 32.7% for humans vs R^2^ = 14.9% for mice in the Gouwens dataset), perhaps in part reflecting the overall degree of greater microglial contamination among the human samples.

### The microglial transcriptomic signature in Patch-seq is reflective of a distinct neuro-inflammatory state

We next sought to identify any distinctive transcriptomic characteristics of microglial contamination in Patch-seq. In particular, we asked whether transcripts concomitantly sampled with neurons reflect particular microglia cell states, such as those observed in disease or in response to injury, or those known to alter neuronal function. For these analyses, we focus on Patch-seq datasets (Berg and Gouwens) collected by the Allen Institute, allowing direct comparison of these transcriptomes to those from dissociated nuclei and cells collected at the same facility using similar methods (see Methods).

To determine the *“transcriptomic signature of Patch-seq microglial contamination”*, we contrasted transcriptomes from neuronal samples with the highest microglial contamination to those with the lowest contamination (see Methods). Numerous genes were overexpressed in Patch-seq neurons with high contamination (Figure 3A, C, human: Supplementary Table 1, mouse: Supplementary Table 2), with 567 genes in human samples and 422 genes in mouse samples (log2 fold-change >2.5; p-value < 0.01). Crucially, this analysis revealed a number of genes that were upregulated in Patch-seq samples with high microglial contamination that are further known to mediate microglial cells’ role in regulating aspects of neuronal excitability, including TNF, IL1B, IL6, and CCL2 (Hikosaka et al., 2022); these example genes are particularly salient as they were not also used to define the microglia contamination score (see Methods).

**Figure 3.**
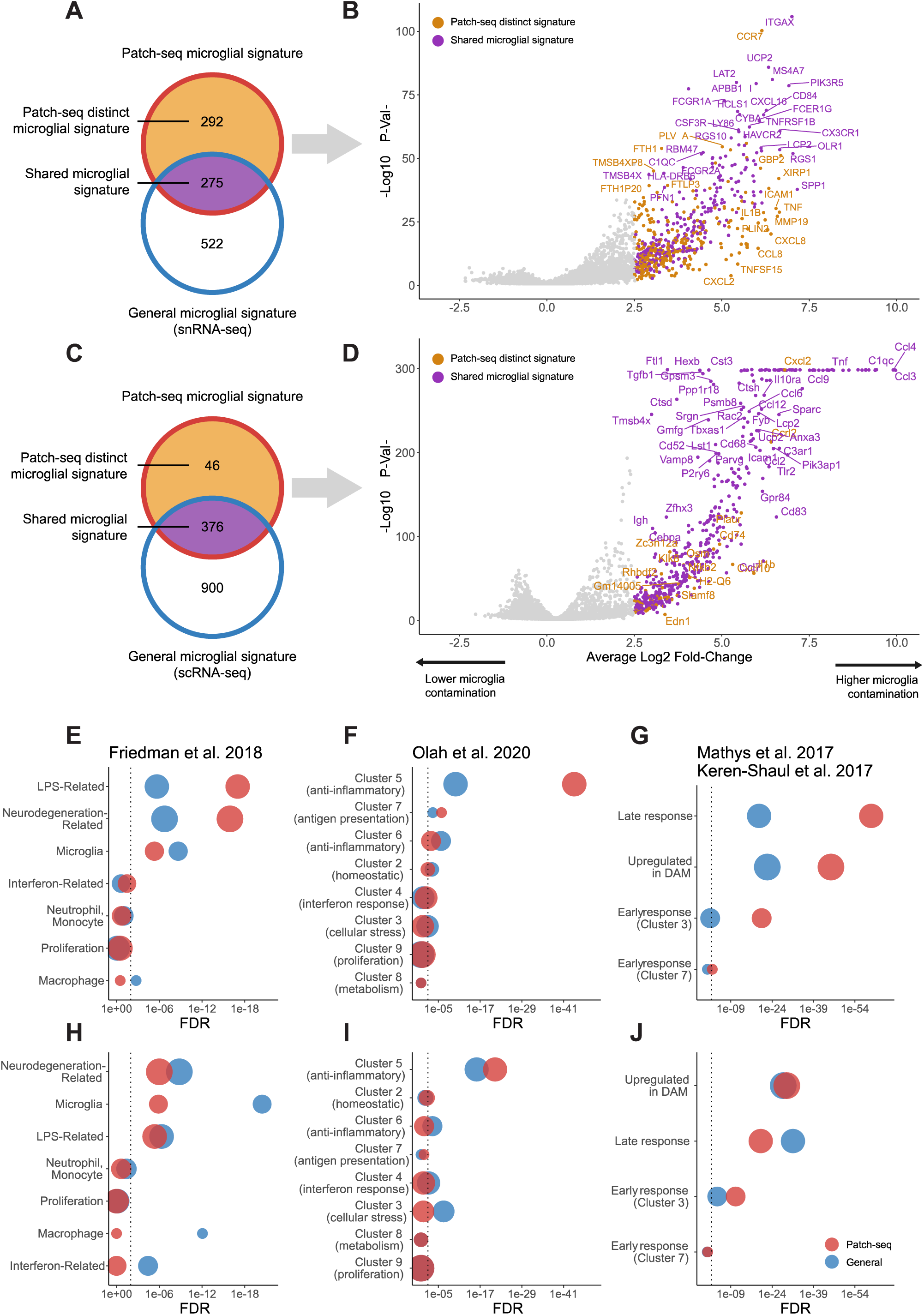
Microglial contamination in Patch-seq reflects a distinct transcriptional signature related to microglia activation. **(A**,**C)** Venn diagrams indicating the number of genes that represent transcriptional signatures of general microglia in (A) human dissociated single-nucleus or (B) mouse single-cell datasets (blue border), transcriptional signature of Patch-seq microglial contamination in human (A) or mouse (B) Patch-seq datasets (red border), genes that are shared between the Patch-seq microglial and general microglia signatures (purple fill), and genes that are distinct to the Patch-seq microglial signature that are not also present in the transcriptional signatures of general microglia (yellow fill). **(B**,**D)** Volcano plots of transcriptional signature of Patch-seq microglial contamination illustrating differentially expressed genes in Patch-seq datasets between human (B) and mouse (D) neuronal samples with high vs. low microglial contamination. Points denote differentially expressed genes (log2 fold-change >2.5; p-value < 0.01), and colors are as in (A, C). Genes in panel (D) with -log10 P Values at 300 indicate significance levels beyond the machine precision limit. **(E-J)** Enrichment analysis of general microglia (blue) and Patch-seq microglia (red) transcriptional signatures (as in A, C) intersected with gene sets of diverse microglial phenotypes and states from multiple data sources (titles in E-G). E-G denotes human microglia signatures and H-J denotes mouse. Dot size reflects the number of genes in each gene set. Dotted line is FDR = 0.05.

Next, we defined a *“general microglia transcriptional signature”*, where we used available gene expression reference profiles from dissociated single-nucleus (human) or single-cell (mouse) RNAseq to identify genes overexpressed in microglia in comparison to neurons. We identified 797 such genes in human and 1276 genes overexpressed in mouse microglia. Lastly, we defined a “*Patch-seq distinct microglial signature”* by identifying genes from the *“transcriptomic signature of Patch-seq microglial contamination”* signature that were not also overexpressed in the *“general microglia transcriptional signature”*. Many genes were identified in the Patch-seq distinct microglial signature, which appeared specific to samples with high levels of microglial contamination in Patch-seq but not dissociated microglia relative to neurons; for example, we saw 292 such genes in humans and 46 such genes in mice (Figure 3B, D). While this analysis might suggest a unique transcriptional signatures in humans relative to mice, there is likely a greater difference between microglial transcripts sampled via Patch-seq in comparison to those sampled via single-nucleus (human) versus single-cell dissociation (mouse), as recent reports have suggested single-nucleus RNAseq is especially limited in the context of microglia-related analyses (Thrupp et al., 2020).

To elaborate on the transcriptional signatures associated with Patch-seq microglial contamination, we performed enrichment analyses of several published reference gene sets that capture diverse microglial states. We compared Patch-seq signatures to our general microglial signatures, to better understand how they may differ. Among human patch-seq samples, we found evidence for an activated, inflammation-relation microglial signature among Patch-seq samples with high microglial contamination. Specifically, the transcriptomic signature of Patch-seq microglial contamination was considerably more enriched for LPS-related (FDR = 1.0 * 10^−17^) and neurodegeneration-related (FDR = 1.2 * 10^−16^) gene signatures defined in Friedman et al. compared to the general microglia transcriptional signature (Figure 3E). Relative to microglia clusters identified in single-cell RNAseq from aged and Alzheimer’s human samples (Olah et al., 2020), the Patch-seq microglial signature was highly enriched for microglia-specific clusters 5, reflecting anti-inflammatory responses (FDR = 8.7 * 10^−45^, Figure 3F). There was also enrichment of microglial late response genes (FDR = 1.2 * 10^−60^) and disease-associated microglia (DAM, FDR = 4.9 * 10^−46^) described by Keren-Shaul et al. and Mathys et al. (Figure 3G). In mouse, we observed few major differences in enrichments between the Patch-seq microglial signature and the general microglial signature (Figure 3H-J), again suggesting fewer differences between microglia sampled during Patch-seq and reference microglial transcriptomes. In summary, these analyses point to an activated, inflammation-related microglial signature among Patch-seq samples with high microglial contamination, in part, which appears distinct from microglial signatures sampled via single-nucleus RNAseq.

### Microglial contamination is associated with alterations in intrinsic electrophysiology

Lastly, we next wanted to assess whether microglial contamination in Patch-seq might be associated with altered excitability of the recorded neuron. We were especially motivated to study this following our observations that microglia contamination is associated with the upregulation of specific transcripts known to causally mediate downstream changes in neuronal excitability (Hikosaka et al., 2022). Because each Patch-seq sampled neuron was first characterized for its intrinsic electrophysiological characteristics prior to mRNA harvesting, we were able to leverage this data to ask how microglial contamination might be associated with cell-to-cell variability in electrophysiological features across neuron subtypes and species.

For the purposes of illustration, we first considered a representative example of two human FREM3 pyramidal cells (Figure 4C), one with low (0.06 microglia contamination score, Figure 4C, blue traces) and one with high microglial contamination (1.00 microglia contamination score, Figure 4C, red traces) but otherwise matched in their overall characteristics (same neurosurgical donor, neuron cell type, similar cortical depths [355 vs 331 um], etc). We observed a number of electrophysiological characteristics that strikingly differed between these two representative cells. For example, the more contaminated cell had a lower input resistance (96 vs 167 MOhms), greater rheobase (110 vs 70 pA), a more depolarized resting membrane potential (−67.8 vs -70.6 mV), and more depolarized minimum voltage following an action potential (i.e., AP trough voltage, -42.8 vs -45.9 mV) than the less contaminated cell, among other differences. We note that these differences, while striking, are within the expected ranges of variability for these cells (Kalmbach et al., 2018; Moradi Chameh et al., 2021) and that the original electrophysiological datasets have undergone strict quality control (Berg et al., 2021). Across human FREM3 pyramidal cells, we also saw that higher microglial contamination scores were associated with lower input resistances (Figure 4A, Pearson’s R = -0.22, p = 0.011) and more depolarized action potential through voltages (Figure 4B, left, Pearson’s R = 0.17, p = 0.047). To help put these associations in context, microglial contamination explained 5.2% of the cell-to-cell variation in input resistance values among human FREM3 pyramidal cells. However, this value is similar to the variance in input resistance explained by the depth of the recorded neuron from the pial surface, 5.1%, noted previously to be a major biological factor distinguishing human superficial pyramidal cells from one another (Berg et al., 2021; Kalmbach et al., 2018; Moradi Chameh et al., 2021).

**Figure 4.**
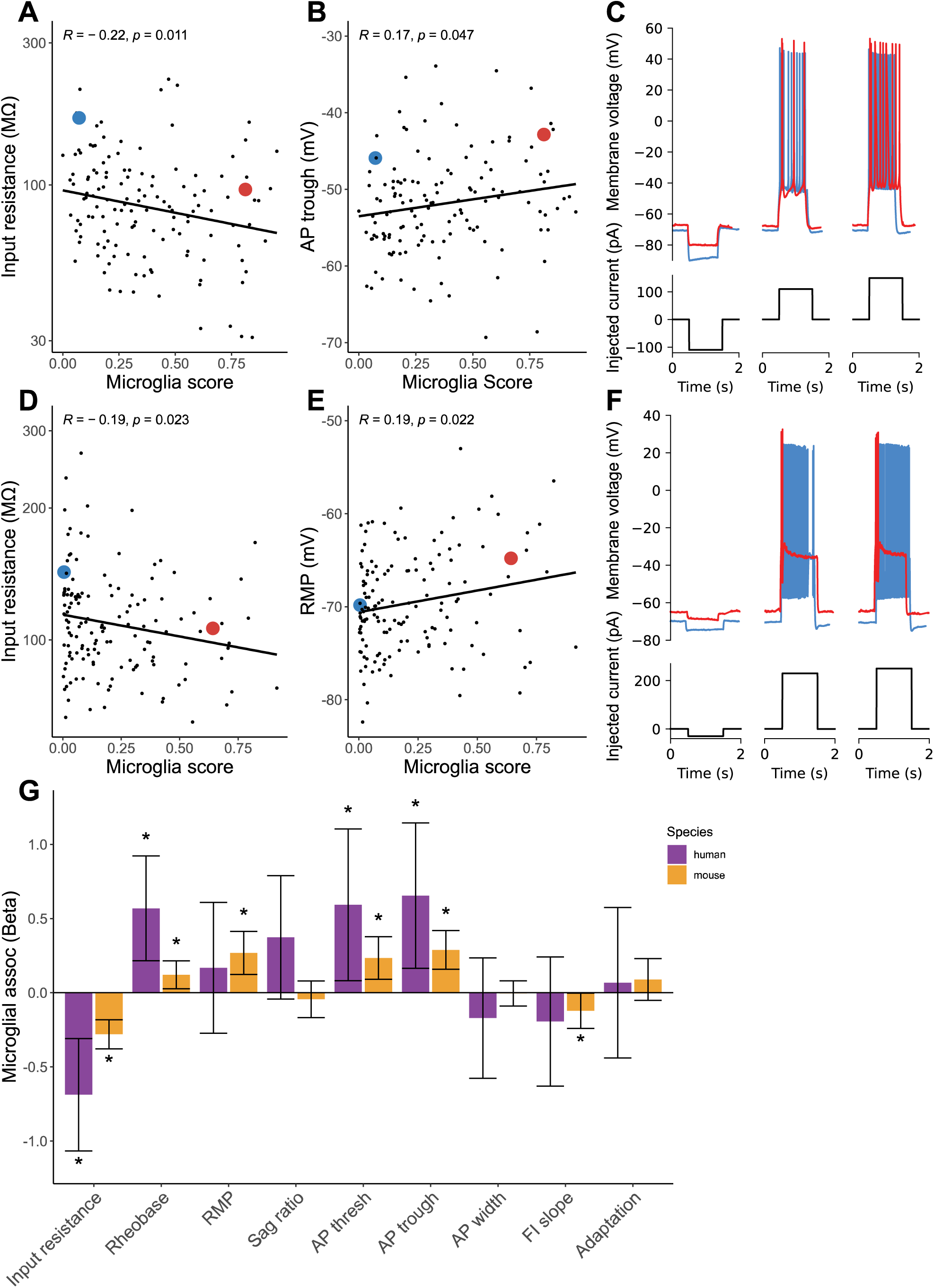
Microglial contamination is associated with altered neuronal intrinsic electrophysiology. **(A, B)** Scatter plots illustrating cellular input resistances (y-axis, A) and action potential trough (y-axis, B) versus microglial contamination scores (x-axis). Each dot reflects one human FREM3 pyramidal cell sampled via Patch-seq. Inset correlations reflect Pearson’s correlations and line indicates best linear fit. Inset blue and red dots reflect two pyramidal cells matched by neurosurgical donor, cell type (FREM3), and cortical depths, but with differing levels of transcriptomically-inferred microglial contamination. **(C)** Membrane voltage traces (top) and corresponding injected currents (bottom) for neurons highlighted in A, reflecting different subthreshold (left) and suprathreshold (middle, right) characteristics of exemplar cells. **(D-F)** Same as (A-C) for mouse Pvalb Sema3e cells from the Gouwens dataset. Resting membrane potentials are shown on y-axis in (E). **(G)** Association between microglial contamination and electrophysiological characteristics, as estimated using a mixed effects model. Bars indicate effect sizes (Beta coefficients) and error bars denote 95% confidence intervals. Asterisks denote beta coefficients where p < 0.05 (ANOVA). Positive (negative) beta coefficients indicate increased microglial contamination is associated with an increase (decrease) in the electrophysiological property. Electrophysiological features have been standardized to unit variance, enabling comparison of beta coefficient effect sizes between species.

As an additional illustrative example, we further considered examples of Pvalb Sema3e cells (reflecting deep layer fast-spiking cells, (Gouwens et al., 2020)), chosen from the same animal but with one example cell expressing low (0.005 microglia contamination score, Figure 4F, blue traces) and the other higher microglial contamination (0.64 microglia contamination score, Figure 4F, red traces). Among these examples and other cells from the same t-type, we observed that cells with higher microglia contamination displayed decreased input resistances (Figure 4D, Pearson’s R = -0.19, p = 0.023) and more depolarized resting membrane potentials (Figure 4E, Pearson’s R = 0.19, p = 0.022), and greater failure in sustaining high frequency firing during sustained depolarizing current injection (Figure 4F)

To quantify the association between microglial contamination and electrophysiological features more systematically, we used a statistical approach to ask how microglial contamination is associated with cell-to-cell variability in electrophysiological features, using mixed effects models to control for cellular and donor/animal identity (see Methods). Among human pyramidal neurons, we found increased microglial contamination associated with multiple electrophysiological features (Figure 4G, statistics in Supplementary Table 3). Most strikingly, we found that increased microglial contamination was associated with significantly decreased input resistances (Beta = -0.68, SE = ±0.19, p = 4.5 * 10^−4^). In addition, a number of suprathreshold features, including action potential threshold and trough voltages, were also significantly associated with microglial contamination (Beta = 0.59, SE = ±0.26, p = 0.024; Beta = 0.65, SE = ±0.24, p = 0.0093). Among GABAergic mouse neurons from the Gouwens dataset (Figure 4G, Supplementary Table 3), we also observed decreased input resistances (Beta = -0.28, SE = ±0.50, p = 2.2 * 10^−8^) and increased action potential threshold and trough voltages (Beta = 0.29, SE = ±0.073, p = 0.0013; Beta = 0.65, SE = ±0.066, p = 1.5 * 10^−5^). We further saw increased microglial contamination associated with more depolarized resting membrane potentials (Beta = 0.26, SE = ±0.074, p = 2.8 * 10^−4^) and a trend towards decreased frequency-current (FI) curve slopes (Beta = -0.12, SE = ±0.060, p = 0.045). We saw similar directions of association between microglia contamination and electrophysiological properties of glutamatergic and GABAergic mouse neurons from the Scala dataset (Supplementary Figure 3, Supplementary Table 3), including decreased input resistances (glutamatergic: Beta = -0.61, SE = ±0.29, p = 0.033, GABAergic: Beta = -0.41, SE = ±0.20, p = 0.049) and increased sag ratios among glutamatergic cells (Beta = 1.0, SE = ±0.31, p = 0.0015). Together, while these statistical associations are correlational, given the temporal nature of Patch-seq where electrophysiological features are recorded prior to mRNA harvest, it is plausible that these associations reflect the direct influence of microglia on neuronal function.

## Discussion

Our analyses of three large-sample human and mouse Patch-seq datasets collected from acute brain slices suggest that microglial off-target contamination is widely present in the transcriptomes of the sampled neurons. A number of technical and biological factors were associated with microglial off-target contamination, including donor-specific factors, particularly in human neurosurgical biopsies, and neuronal cell type identity. The transcriptomic signature of microglial off-target contamination in Patch-seq appears indicative of a distinct activated and inflammation-related cell state. Critically, microglial contamination is further associated with altered neuronal electrophysiological features, including lowered input resistances and increased action potential thresholds, suggesting provocatively that microglia are shaping intrinsic properties of the recorded neurons.

While the presence of off-target cellular contamination in neuronal Patch-seq datasets has been reported previously, we were somewhat surprised by how prevalent it appeared in the datasets analyzed here. Moreover, we were concerned that microglial contamination appeared to be a major contributor to unbiased clustering of Patch-seq derived neuronal transcriptomes when not excluded by prefiltering. Microglial off-target contamination differed considerably in samples collected from different donors; in human datasets specifically, this might reflect the inherent challenges in obtaining biopsies from neurosurgical tissue, which we and others have suggested contributes to technical variability in efforts to systematically characterize human neurons in brain slices (Moradi Chameh et al., 2021). However, we note that animal identity was among the top two factors related to microglial contamination among both of the mouse Patch-seq datasets analyzed, suggesting that it may be an important proxy for tissue health or slice preparation quality that might differ systematically across animals. Similarly, our analyses of different levels of observed microglial contamination among mouse neocortical neurons might be reflective of differential vulnerability of these cell types to the influence of microglia *in vivo*. For example, we found that Pvalb Vipr2 cells, reflective of chandelier cells, displayed the highest levels of microglial contamination among all tested mouse GABAergic neuronal types across two datasets sampling different neocortical brain regions; intriguingly a recent report suggests that microglia are key regulators of chandelier cell axonal arborization and synapse formation (Gallo et al., 2022).

The transcriptional signature associated with the involvement of microglia in Patch-seq appeared indicative of activated microglia. This inference is based on the overexpression of key hallmark genes, including TNF and CCL2, and that the Patch-seq related microglial transcriptional signature shares similarities with other microglia signatures related to LPS-, neurodegeneration, and disease associated microglia (Friedman et al., 2018; Keren-Shaul et al., 2017; Mathys et al., 2017; Olah et al., 2020). In addition, we note that we saw considerably more microglia-related genes distinct to human versus mouse Patch-seq datasets. While this might be reflective of bona fide species differences, one simple explanation for this finding may be related to our usage of single-nucleus RNAseq to provide reference profiles of microglia from humans but single-cell RNAseq from mice. Recently, it has been shown that single-nucleus RNAseq, when used to profile human microglia, are more prone to missing important transcripts related to microglial proliferation and other disease related processes (Thrupp et al., 2020). Such transcripts are possibly expressed in distal cellular processes and are likely to be especially present in microglia inadvertently sampled during Patch-seq.

A key question our findings raise is how contact between microglia with neurons, either directly or indirectly, might contribute to altered cellular electrophysiology. We hypothesize two possible explanations that consider the order of events in Patch-seq, where cellular electrophysiology is characterized before mRNA is harvested and sequenced (Lipovsek et al., 2021). First, we hypothesize that microglia might be directly interacting with the characterized neuron, for example, via physical interactions (Cserép et al., 2020) or through signaling molecules like TNF that induce downstream changes in cellular excitability through phosphatase signaling and calcium-activated potassium channels (Yamamoto et al., 2019; Yamawaki et al., 2022). A second alternative hypothesis is that microglial processes might be chemotaxing near neurons that have been otherwise altered or damaged, for example by the slice preparation procedure, but such neuron-microglia interactions might not be directly causing alterations to cellular electrophysiology. Because of the observational nature of our study, we cannot directly reconcile these competing explanations as doing so would likely require further experiments that directly perturb microglia; nevertheless, our findings are consistent with a growing body of work indicating that microglia directly regulate aspects of neuron excitability (Badimon et al., 2020; Cserép et al., 2020; Hikosaka et al., 2022).

Our study has a number of limitations. First, we note that our analyses rely on our ability to reliably infer the presence of microglial processes in proximity to the characterized neuron using transcriptomics (Tripathy et al., 2018). We feel reasonably confident in this inference as it is unlikely for neuronal cells to endogenously express tens of microglial specific markers and there is a reasonable concordance between microglial mRNA expression and microglia marker protein expression assayed via immunohistochemistry. Second, we note that our analyses are observational in nature, relying on the natural variability inherent to different experiments. As such, it is difficult to conclude cause from effect in our study, for example, whether microglia-neuronal interactions directly contribute to altered intrinsic electrophysiology. Lastly, we were limited in our analyses to the data fields provided by the original study authors. In some cases, this precluded the study of additional, potentially causal factors related to microglial contamination and proliferation, such as the relative timing of neuronal recordings in relation to when acute slices were prepared and additional human donor-specific fields, including factors related to the quality of neurosurgical resection or genetic variants related to microglial function (Felsky et al., 2019).

Our study raises a number of questions on the nature of microglia-neuron interactions. First and foremost, we need to better understand microglial activation in the context of experimental work making use of acute brain slices. What contributes to increased microglial activation among slices from some donors and animals, but not others? And can unwanted microglial activation be minimized, for example by incorporating known strategies for inhibiting or depleting microglia, like pre-incubating slices with the CSF1R inhibitor, PLX3397 (Yamamoto et al., 2019; Yamawaki et al., 2022)? Second, what is drawing microglia to the surrounding microenvironment of neurons undergoing patch-clamp electrophysiology? Are microglial processes chemotaxing to the neuron’s soma (as opposed to synapses or dendrites)? Is the presence of ATP or other known chemoattractants in or near the patch pipette attracting microglia (Madry et al., 2018)? Are unhealthy neurons damaged by slice preparation or hypoxia releasing pro-inflammatory cytokines that attract microglia? Lastly, and most importantly, to what extent are the microglia-neuronal associations reported here, in the context of acute brain slices, reflective of such interactions *in vivo*? These questions highlight an important potential role for microglia in shaping neuronal excitability and reflect a number of questions for future investigation.

## Acknowledgements

We acknowledge the original study authors for graciously making their datasets publicly available which enabled this work. We thank Jeremy Miller, Brian Lee, Rachel Dalley, and members of the Tripathy laboratory for comments and feedback. This study was supported by the University of Toronto Institute for Medical Science Undergraduate Research Opportunity Program for Epilepsy or Muscular Dystrophy Research, CAMH Discovery Fund, Krembil Foundation, Kavli Foundation, McLaughlin Foundation, Natural Sciences and Engineering Research Council of Canada (RGPIN-2020-05834 and DGECR-2020-00048), Canadian Institutes of Health Research (NGN-171423, NGN-435539, and PJT-175254), the Simons Foundation for Autism Research, and National Science Foundation (NeuroNex Grant NSF2015276).

## Supplementary

**Supplementary Figure 1.**
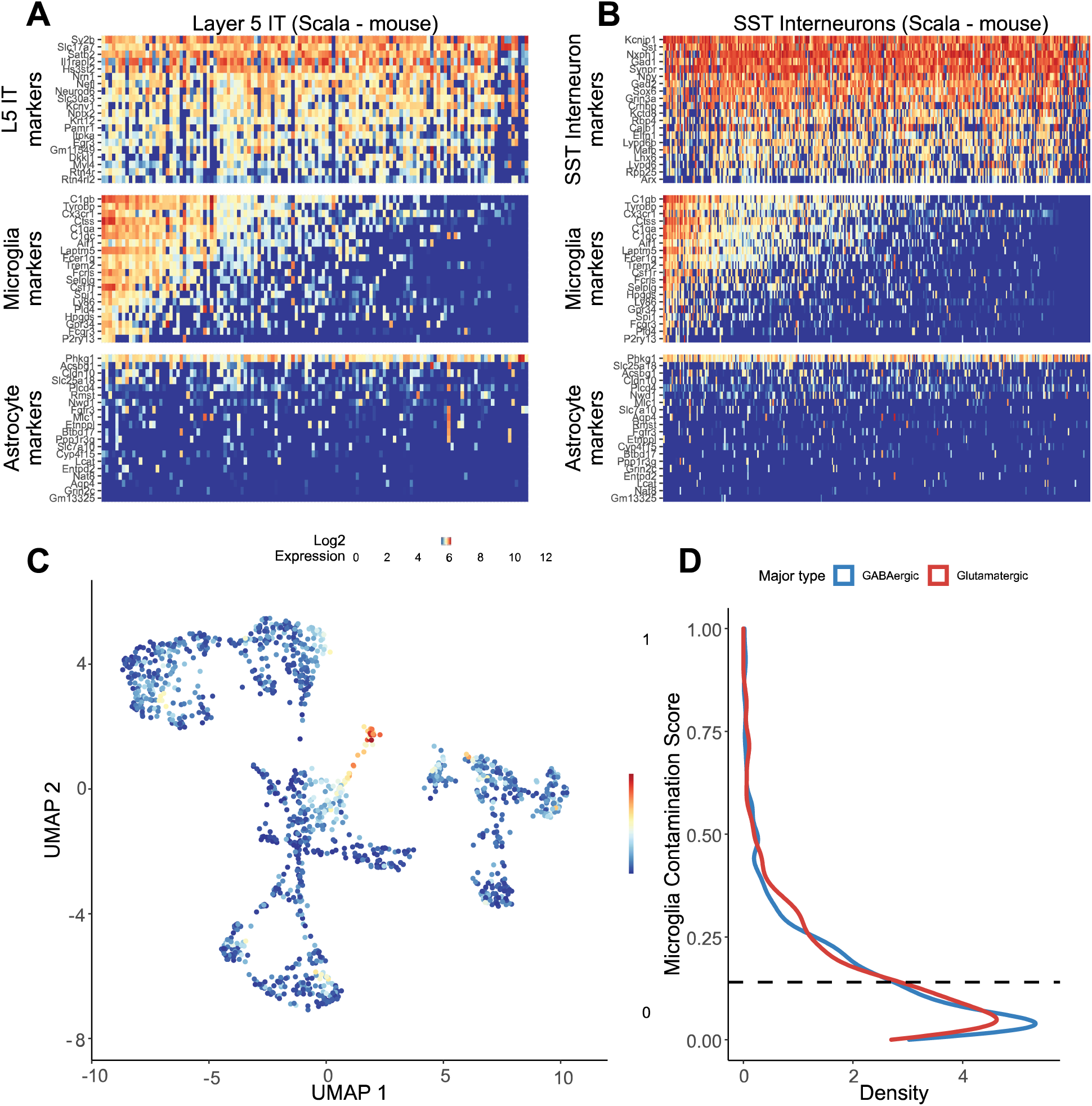
Patch-seq transcriptomes of glutamatergic and GABAergic mouse neurons from the Scala dataset express off-target expression of microglial marker genes. **(A**,**B)** Gene expression profiles for Layer 5 Intratelencephalic neurons (A) or SST interneurons (B) for various cell type-specific markers. Each row represents a cell type-specific marker gene and columns represent individual neurons, ordered from left to right by decreasing microglial contamination score. **(C)** Low-dimensional visualization of transcriptomes from neuronal Patch-seq samples clustered by most variable gene expression and color-coded by microglial contamination score. **(D)** Distribution density of microglial contamination scores for GABAergic (blue) or glutamatergic (red) neurons. Dashed lines indicate population mean.

**Supplementary Figure 2.**
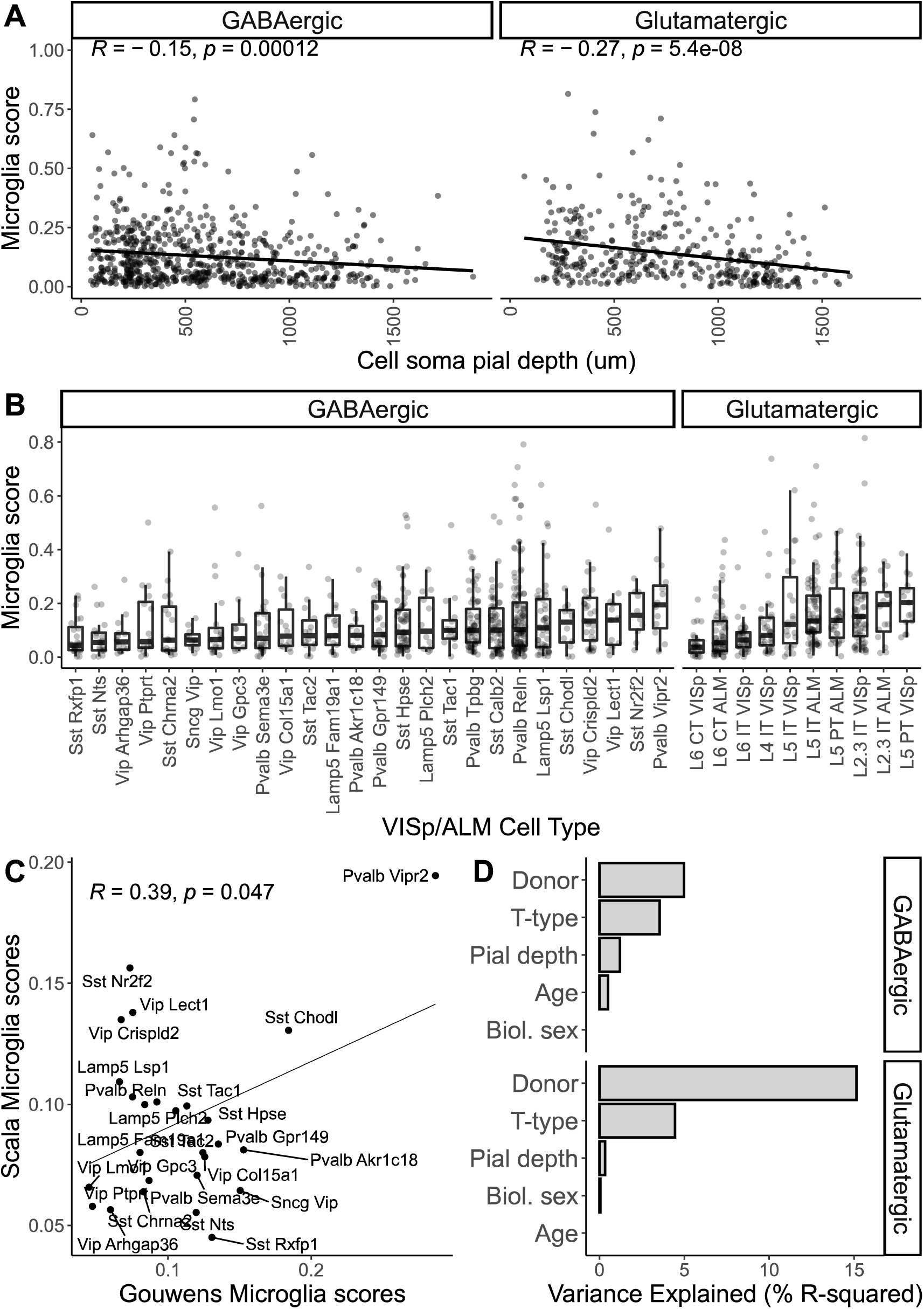
Associations between cell soma pial depth, cell type, and other factors with off-target microglial expression in Patch-seq samples from the Scala mouse dataset. **(A)** Associations between microglia contamination scores (y-axis) estimated from mouse GABAergic and Glutamatergic neuron Patch-seq samples with recorded depths of cell soma from the pial surface (x-axis). Inset correlations reflect Pearson’s correlations and line indicates best linear fit. **(B)** Associations between microglia contamination scores (y-axis) and cell type identity summarized at the t-type or cluster level (mapped to the same VISp/ALM cell type atlas used in the Gouwens dataset). **(C)** Comparison of cell type-specific Microglia scores among GABAergic cells from the Scala dataset (y-axis) and Gouwens dataset (x-axis). Each dot reflects median Microglia contamination scores for one transcriptomically-defined cell type. Inset correlation reflects Pearson’s correlation and line indicates best linear fit. **(D)** Estimated percent variance explained (R-squared) in microglial contamination scores among GABAergic (top) and Glutamatergic (bottom) samples by various factors, including donor/animal identity, neuronal cell type identity (t-type), age, sex, and sample depth from pial surface (Pial depth).

**Supplementary Figure 3.**
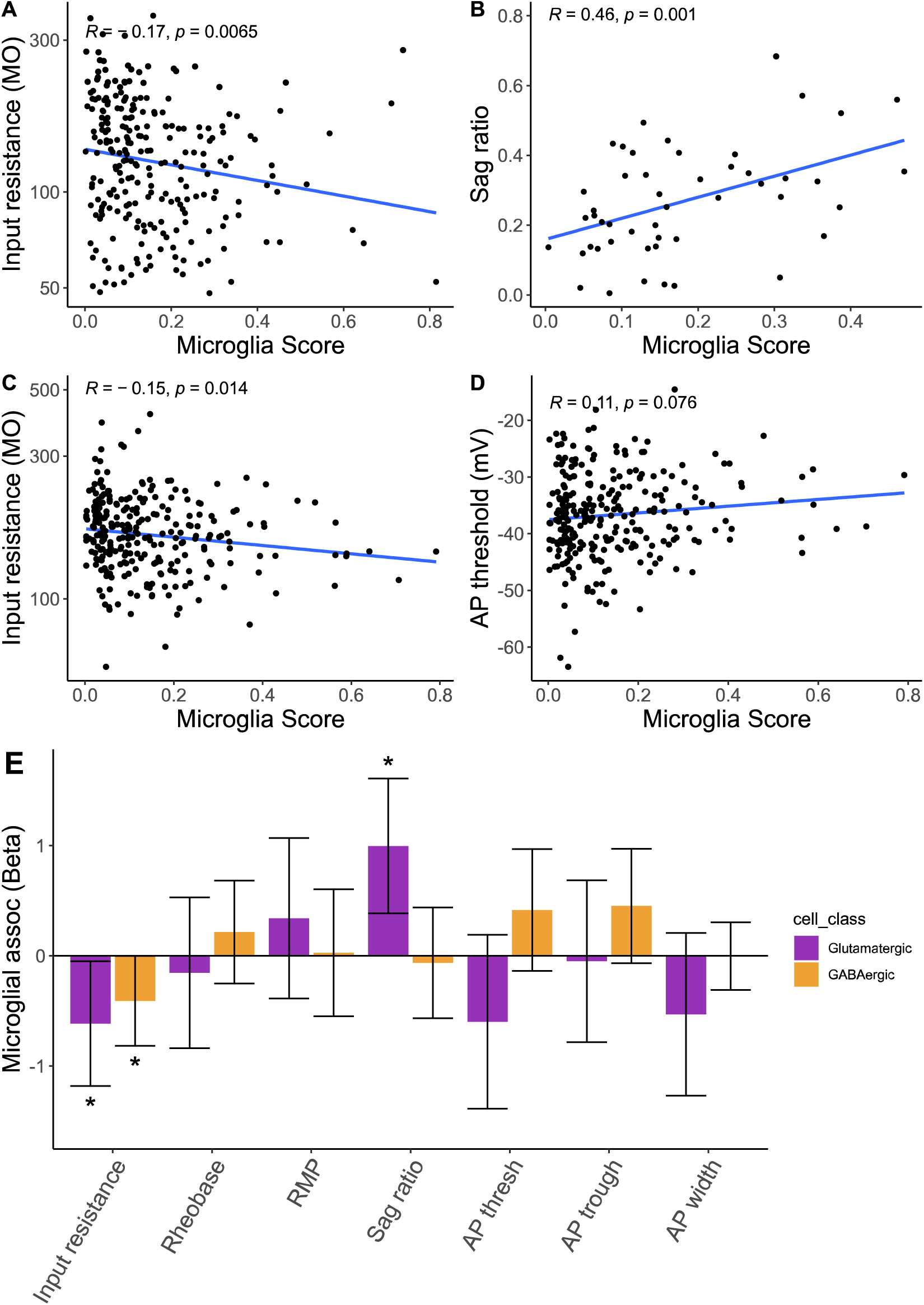
Associations between microglial contamination and neuronal intrinsic electrophysiology in the Scala mouse dataset. **(A-D)** Scatter plots illustrating electrophysiological features (y-axis) versus microglial contamination scores (x-axis). Lines reflect lines of best fit and inset correlation values denote Pearson’s correlations. (A) Input resistance values from intra-telencephalic pyramidal cells; (B) Sag ratio values from extra-telencephalic pyramidal cells; (C, D) Input resistance values (C) and action potential threshold values (D) from Pvalb interneurons. **(E)** Association between microglial contamination and electrophysiological characteristics, as estimated using a mixed effects model. Bars indicate effect sizes (Beta coefficients) and error bars denote 95% confidence intervals. Asterisks denote beta coefficients where p < 0.05 (ANOVA). Negative (positive) beta coefficients indicate increased microglial contamination is associated with a decrease (increase) in the electrophysiological property. Electrophysiological features have been standardized to unit variance, enabling comparison of beta coefficient effect sizes between species.

**Supplementary Table 1. *Human Patch-seq differential expression between cells with high vs. low microglial contamination***

***https://github.com/keon-arbabi/patch-seq-microglia/blob/main/output/supplementary_tables/supp_table_1.csv***

**Supplementary Table 2. *Mouse Patch-seq differential expression between cells with high vs. low microglial contamination***

***https://github.com/keon-arbabi/patch-seq-microglia/blob/main/output/supplementary_tables/supp_table_2.csv***

**Supplementary Table 3. *Effect sizes for associations between microglial contamination and electrophysiological features from mixed effects models***

***https://github.com/keon-arbabi/patch-seq-microglia/blob/main/output/supplementary_tables/supp_table_3.xlsx***

### Available code

***https://github.com/keon-arbabi/patch-seq-microglia.git***

